# Microbiological safety of industrially reared insects for food: Identification of bacterial endospores and targeted detection of foodborne viruses

**DOI:** 10.1101/2020.04.22.055236

**Authors:** D. Vandeweyer, B. Lievens, L. Van Campenhout

## Abstract

Edible insects are characterised by high microbial numbers of which the bacterial endospores fraction can survive (thermal) processing. It is unknown, however, which bacterial species occur as endospore in edible insects and what impact they have on food safety. Additionally, edible insects have never been explored for the presence of foodborne viruses so far. In this study, we found that the bacterial endospore fraction in a collection of mealworm and cricket samples obtained from commercial insect producers can comprise a large amount of *Bacillus cereus* group members that can pose insect or human health risks. Monitoring and effective mitigation of these risks are urged. By contrast, none of the foodborne viruses hepatitis A virus, hepatitis E virus and norovirus genogroup II were detected in the sample collection. Therefore, food safety risks originating from these viral pathogens can be concluded to be low.

## Introduction

As from 2014, edible insects and insect-derived food products started to enter the European markets. As a result, risk assessments and food safety research regarding this new food matrix were initiated, e.g. by EFSA^1^. In the past five years, many basic microbiological food safety questions were formulated and investigated, as reviewed by Garofalo *et al.* (2019)^2^. In summary, freshly reared insects may be highly contaminated with diverse microorganisms, but these high numbers can easily be reduced, e.g. by applying a mild heat treatment. This was demonstrated both for fungi and bacteria for yellow mealworms (*Tenebrio molitor*)^3^, lesser mealworms (*Alphitobius diaperinus*)^4^ and tropical house crickets (*Gryllodes sigillatus*)^5^. However, it was repeatedly observed that a fraction of the bacteria, i.e. the endospores, can survive those mild heat treatments^3,6^. *Bacillus* and *Clostridium* are two relevant bacterial genera that are able to produce endospores. Since several members of these genera are food pathogens (e.g. *B. cereus, C. perfringens, C. botulinum*), the survival of endospores in the insect matrix may pose food safety risks. DNA-based studies have already identified members of both genera in edible insects and insect products produced in Europe^7–11^, but it is generally not clear which specific species were found and whether or not they were present in spore form. Consequently, the first goal of this research (experiment 1) was to identify which bacterial endospores can be present in the two edible insect species reared mostly for human consumption nowadays: the yellow mealworm (*Tenebrio molitor*) and the house cricket (*Acheta domesticus*). Specific attention was given to *B. cereus*-related isolates, which were further characterised by exploring specific genes with qPCR.

Another aspect that has not been explored so far is the presence of pathogenic foodborne viruses in insects for food. The possibility that foodborne viruses such as hepatitis A virus (HAV), hepatitis E virus (HEV) or norovirus (NoV) can be present in insects or derived foodstuffs has been proposed by EFSA already in 2015^1^. The three said viruses are the most commonly transmitted viruses through foodstuffs^12^ and are relevant to be investigated for insects and insect-based food matrices. This research was the first to elucidate whether the selected human foodborne viruses HAV, HEV and NoV genogroup II (GII, responsible for the majority of norovirus cases^13^) are present in mealworm and cricket species, and if so, in what quantity (experiment 2).

When studying the microbiota of insects, it is very plausible that the organisms present in their gut and on their exoskeleton differ between insects cultivated in the laboratory and the same species reared at industrial scale, where the hygiene levels may be lower and hence the house flora different. Since the aim of this work was to study bacterial endospores and viruses with respect to safety for the consumer, insect samples obtained from several commercial insect producers were studied and multiple batches per producer were included in the experimental set-up.

## Results and discussion

### Bacterial endospore counts

In this study, the samples investigated were first subjected to bacterial endospore counts and, as a comparison, total viable counts (Table 1). The obtained values for total viable aerobic (between 7.5 and 8.7 log cfu/g) and anaerobic (7.4 to 8.7 log cfu/g) counts are similar to the typically high counts for raw edible insects reported earlier^2,14^. Additionally, the values for the aerobic and anaerobic counts are highly comparable. This may suggest that a large fraction of both counts exists of facultative anaerobic species. When comparing both insect species investigated, it is clear that the total viable counts for house crickets are higher than those for yellow mealworms. This confirms what was proposed earlier^15^, being that microbiological findings of particular insect species should not be generalised for all edible insects, as nevertheless is done in literature occasionally. For the total viable counts, small and insignificant differences were observed between samples of the same species. For bacterial endospore counts, on the other hand, larger and significant variation was observed between samples from the same insect species, compared to the total viable counts. This variation may result from differences in (hygiene in) the rearing environment and/or feed, and is commonplace for insects^2,9,11^. Again, anaerobic and aerobic endospore counts displayed comparable trends (Table 1).

**Table 1.** Total and endospore counts of the samples investigated. Data are the mean values of three replicates ± standard deviation.

| Insect species | Rearing company | Batch | Microbial counts (log cfu/g) |  |  |  |
| --- | --- | --- | --- | --- | --- | --- |
|  |  |  | Total viable aerobic count | Total viable anaerobic count | Aerobic bacterial endospores | Anaerobic bacterial endospores |
| Yellow mealworm | 1 | 1 | $7.5 \pm 0.1^a$ | $7.4 \pm 0.2^a$ | $2.6 \pm 0.5^a$ | N.D.<sup>*</sup> |
| | | 2 | $8.0 \pm 0.0^a$ | $7.7 \pm 0.6^a$ | $2.3 \pm 0.0^a$ | $1.6 \pm 0.4^a$ |
| | 2 | 1 | $8.1 \pm 0.1^a$ | $8.1 \pm 0.0^a$ | $2.6 \pm 0.9^a$ | $2.6 \pm 0.9^{a,b}$ |
| | | 2 | $7.7 \pm 0.5^a$ | $7.8 \pm 0.5^a$ | $3.4 \pm 0.5^b$ | $3.7 \pm 0.6^b$ |
| House cricket | 3 | 1 | $8.7 \pm 0.3^a$ | $8.5 \pm 0.2^a$ | $3.1 \pm 0.1^a$ | $3.3 \pm 0.5^{a,b}$ |
| | | 2 | $8.3 \pm 0.0^a$ | $8.3 \pm 0.1^a$ | $3.4 \pm 0.5^a$ | $2.2 \pm 0.7^a$ |
| | 4 | 1 | $8.7 \pm 0.3^a$ | $8.7 \pm 0.3^a$ | $4.2 \pm 0.5^b$ | $3.9 \pm 0.9^c$ |
| | | 2 | $7.8 \pm 0.0^a$ | $7.8 \pm 0.0^a$ | $3.9 \pm 0.4^b$ | $4.0 \pm 0.9^{b,c}$ |

<sup>\*</sup>N.D. = not determined.
<sup>a,b,c</sup>Mean counts from different samples per insect species with the same superscript within the same column are not statistically different ( $p > 0.05$ ).

Currently in the insect sector, edible insects are typically subjected to a heat treatment prior to processing and consumption, such as boiling and/or oven drying. It has been demonstrated^3,5,6^ that bacterial endospores can survive these treatments and remain present and viable in insects for human consumption. The highest endospore count observed in this study was 4.2 log cfu/g (for a house cricket sample). While no European microbiological criteria exist specifically for edible insects, this count does surpass the lower action limit for *Bacillus cereus* in edible insects as published by the Belgian Federal Agency for the Safety of the Food Chain^16^. In the case that a large fraction of these spores found would consist of *B. cereus*, this would pose a health risk for human consumption. For clostridia, no criteria or action limits exist, but it is advised to pay attention to these potential pathogens as well.

### Identification of bacterial endospores

Microbiological health risks of food products are determined not only by population densities of microbial (sub)groups, but also by the particular species present in the foodstuff. In this study, we focussed on bacterial endospores, which are resistant against several treatments that are harmful for vegetative cells. Pathogenic bacteria present in their spore form are of major concern for edible insects.

In total, 142 endospore-forming isolates were collected from the bacterial endospore count plates obtained from yellow mealworm and house cricket samples. These isolates were present as endospore in the insect matrix and survived the pasteurisation treatment, germinated and grew on Plate Count Agar. The isolates were assigned to four different genera of endospore-forming bacteria, including *Bacillus, Lysinibacillus, Brevibacillus* and *Clostridium*. Based on further identification, the *Bacillus* genus was further divided into *Bacillus cereus* group (also *B. cereus* sensu lato (s.l.)) and *Bacillus* (non-cereus) (Figure 1A; Supplementary Table 1). Of the 50 isolates obtained from yellow mealworms, 20 (i.e. 40%) were identified as a member of the *Bacillus cereus* group. For house crickets, even 79 out of 92 isolates (86%) were identified as *B. cereus* s.l. This may indicate that house crickets are highly susceptible to colonisation with *B. cereus* group members.

**Figure 1.**
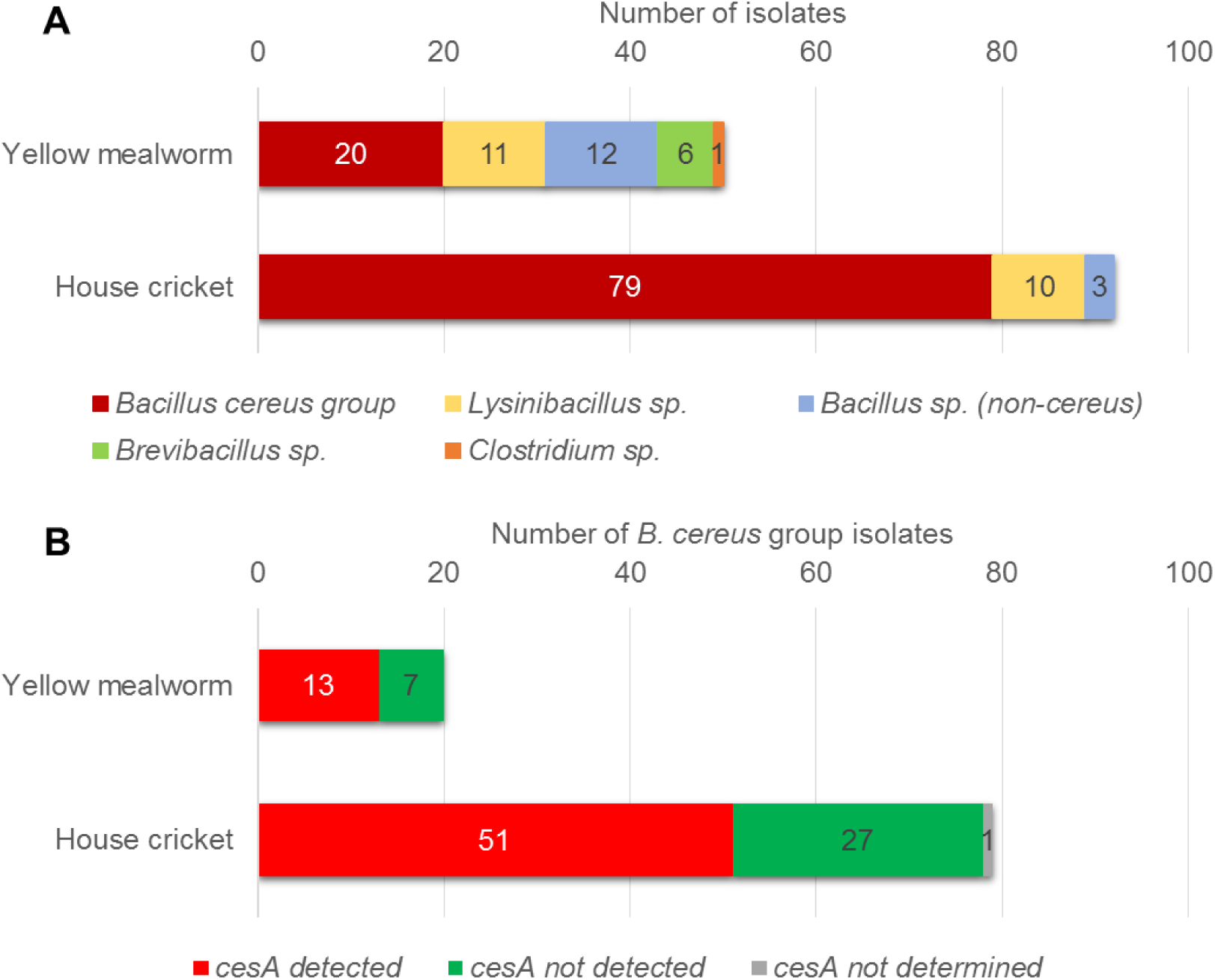
(1A) Number of bacterial endospore isolates from yellow mealworms and house crickets corresponding to the bacterial genera identified by 16 S rRNA gene sequencing and (1B) presence of the *cesA* gene, *in B. cereus* group isolates, as determined by qPCR.

The *B. cereus* group consists, among a few recently described species, of the closely related bacteria *B. cereus* sensu stricto, *B. thuringiensis, B. weihenstephanensis, B. wiedmannii, B. anthracis, B. mycoides, B. pseudomycoides, B. cytotoxicus*, and *B. toyonensis*^17–20^. Most isolates assigned to the *B. cereus* group were identified as *B. cereus* s.s. or *B. paramycoides* (proposed novel member^21^). Due to the close genetic relationships within the *B. cereus* group, however, it is hard to distinguish between the different species compiled in this group based on the 16 S rRNA gene^20^. Several species of the *B. cereus* group are relevant with respect to human health, agriculture or food safety. In the case of edible insects, presence of the well-known pathogen *B. cereus* s.s. may pose a severe food safety risk for consumers, but also some *B. weihenstephanensis* and *B. cytotoxicus* strains were reported as potential human (foodborne) pathogens^22,23^. For insect rearing, *B. thuringiensis* may be harmful because of its insecticidal properties^24^. Whatever *B. cereus* group species were present in the insects, the *B. cereus* group poses an important threat for the edible insect sector and should be monitored. Also, given the fact that the *B. cereus* group contains psychrotrophic strains^19,25^, chilled preservation of heat-treated insects can still allow the growth (and toxin production) of food pathogens such as *B. cereus* s.s.

A few previous studies on the microbiota harboured by edible insects also report members of the *B. cereus* group. For example, *B. cereus* s.s. and *B. weihenstephanensis* were detected by PCR-DGGE (Denaturing Gradient Gel Electrophoresis) in edible processed insects bought in Belgium and the Netherlands^10^. In another study^11^, processed edible insects were described to contain up to 6.6 log cfu/g presumptive *B. cereus*, of which *B. cytotoxicus, B. cereus* s.s. and *B. thuringiensis* isolates could be identified using sequencing and biomolecular identification.

The second most observed endospore-forming bacteria were members of the genus *Lysinibacillus* (21 out of 142 isolates, or 15%; Figure 1A) and the majority was identified as *L. fusiformis* (Supplementary Table 1). This spore former, renamed from *Bacillus fusiformis*^26^, rarely acts as a human pathogen, but especially its relative *L. sphaericus* has been reported as a potent insect pathogen. *L. sphaericus* preferably targets mosquitos, but activity against other insects species has been observed as well^27^. Also previously^11^, *Lysinibacillus* sp. was detected in edible insects.

Besides *B. cereus* members, also other, non-cereus *Bacillus* spp. were detected (16/142 isolates, 11%; Figure 1A). The genus *Bacillus* has regularly been found in edible insects^6,10,11,28,29^, but it has rarely been described at species level. According to the identification results (Supplementary Table 1), non-cereus *Bacillus* isolates may correspond to several species including *B. pumilus, B. altitudinis, B. siamensis, B. vallismortis* and *B. subtilis*. The latter two species are closely related and, comparably to the *B. cereus* group, they are members of the *B. subtilis* group^30^. As was also the case for the *B. cereus* group, the 16 S rRNA gene does not allow for species-level identification for *B. subtilis* group members^30^. *B. subtilis* and its relatives are generally not considered as human pathogens, but may act as spoilage organisms instead^31^. They are also frequently described as beneficial for plants and/or animals e.g. as biocontrol organism or probiotic^32,33^, which may pose opportunities for industrial valorisation, also in the insect sector.

Six isolates (3%; Figure 1A) were ascribed to the genus *Brevibacillus* and identified as either *B. laterosporus* or *B. halotolerans* (Supplementary Table 1). They were only obtained from yellow mealworms and not from house crickets. In previous research^7^, a *Brevibacillus* sp. was detected by Illumina sequencing in yellow mealworms, in abundances up to 28%. *B. laterosporus* has been reported as insect pathogen^34^. Together with *B. thuringiensis*, a member of the *B. cereus* group, and *Lysinibacillus sphaericus*, a realistic risk for edible insect rearing exists when these entomopathogenic bacteria are present. On the other hand, *B. laterosporus* has also been described as a broad-spectrum antimicrobial species against phytopathogenic bacteria and fungi^34^, which may involve agricultural benefits.

While both aerobic and anaerobic bacterial endospore counts were performed and isolates were picked randomly from all plates from all samples, it is striking that almost solely aerobic and/or facultative anaerobic organisms were identified. (Strict) anaerobic spore formers such as specific *Clostridium* spp. were encountered in edible insects before, including yellow mealworms and house crickets^7–9,35^. In this study, however, only one isolated was assigned to the genus *Clostridium*, yet, with a low sequence identity (69.4 %, Supplementary Table 1). The genus *Clostridium* is of concern regarding food safety, since it contains the food pathogens *C. perfringens* and *C. botulinum*^36^. All previous observations of (possible) anaerobic spore-forming bacteria were, however, based on culture-independent methods. The random selection of isolates and/or the strict anaerobic conditions necessary for cultivating certain *Clostridium* species may have impeded to obtain anaerobic species that might have been present in the samples investigated. Accordingly, the specific isolation and identification of anaerobic bacterial spore formers would form an interesting addition to this research.

### qPCR detection of B. cereus-related genes

It has been demonstrated that the 16 S rRNA gene is extremely similar (97.34 to 100% inter-species similarity) between all *B. cereus* group members ^19,37^. The 16 S rRNA gene-based identification method applied in this study was therefore complemented by studying additional genetic markers to further characterise and discriminate the *B. cereus* s.l. isolates. Firstly, all *B. cereus* s.l. were subjected to a qPCR assay targeting the *panC* gene encoding a pantothenate-β-alanine ligase^38^. The presence of this gene was used to confirm the assignment of the isolates to the *B. cereus* group, since *panC* genotyping is a typical method to group *B. cereus* s.l. species based on phenotypic characteristics^39^. Next, by targeting the *cesA* gene^40^, each isolate was assessed for plasmid presence encoding for the production of the heat-resistant cereulide toxin, the most severe food safety risk^41,42^. This allows to evaluate the possibility for each isolate to act as a human pathogen of the emetic type, regardless of its taxonomic classification. Additionally, to date, this virulence plasmid was only encountered in *B. cereus* s.s. and *B. weihenstephanensis*, while it was reported not to be present in a large collection of *B. thuringiensis, B. mycoides* and *B. pseudomycoides* strains^17,42^. In the (edible) insect sector, the distinction between foodborne pathogen (*B. cereus* s.s. or *B. weihenstephanensis*) and insect-borne pathogen (*B. thuringiensis*) is of major interest, and presence or absence of the *cesA* gene provides additional information in this regard.

Of the 98 *B. cereus* group isolates assessed for the *panC* gene, only three were found to be negative. Consequently, the identification of 95 *B. cereus* group isolates was considered reliable. Next, in 64 of the 98 isolates (65 %), the *cesA* gene was detected (Figure 1B). This indicates that those isolates may be able to produce the cereulide toxin under suitable conditions. Influencing factors are for instance temperature, the food matrix and the number of *B. cereus* cells present^43,44^. The amount of *B. cereus* required to be present for toxin production to take place is typically reported as 3 to 5 log cfu/g^17^. *B. cereus* counts were not assessed in this study, but in literature, presumptive *B. cereus* were already detected up to 6.6 log cfu/g in processed edible mole crickets^11^.

The presence of the cereulide plasmid in *B. cereus* in its endospore form poses an increased food safety risk. The number of bacterial endospores in the samples investigated was maximally 4.2 log cfu/g and did not relate solely to *B. cereus* s.l. Despite this observation, a risk is still present because after a heat treatment (which is survived by the spores and can also even activate them), the endospores can germinate in the insect matrix, and in absence of competition, multiply and produce cereulide. A second heating step will not be sufficient to destroy the toxin and thus the risk. Consequently, the exact conditions for and the extent of cereulide production that can occur in edible insects and derived food products is a future research need.

### Presence of foodborne viruses

Three different RT-qPCR assays were optimised to detect and quantify the RNA viruses HAV, HEV and NoV GII in insect RNA extracts and assessed for their reliability. The addition of an Internal Amplification Control (IAC) to the RT-qPCR assay has proven to be a robust control in virus analysis in food matrices^45^ and has become common practice^46,47^. Similarly, in this study, IACs were included in all samples investigated and were systematically recovered in the three assays (Ct values ± 35), while negative IAC controls showed no amplification after 40 cycles, proving the reactions to be successful. In order to quantify the viral titer, standard curves were constructed and showed to be of good quality, according to their calculated parameters (Table 2). Based on the recovery of the reference virus cDNA fragments, the detection limit for each assay was set at 100 copies/µl undiluted RNA extract. Also the consistent amplification of positive controls (Ct values ± 31) and absence of qPCR signal for negative controls contributed to the conclusion that the RT-qPCR assays were reliable.

**Table 2.** Standard curve parameters as calculated by the qPCR software.

| Target | R<sup>2</sup> | Slope | Efficiency (%) |
| --- | --- | --- | --- |
| HAV | 0.998 | -3.737 | 85.192 |
| HEV | 0.999 | -3.287 | 101.487 |
| NoV GII | 0.999 | -3.625 | 88.739 |

In none of the samples any of the viruses HAV, HEV or NoV GII could be detected, meaning that the viruses were either not present or the samples contained less than 100 viral RNA copies/µl extract. Consequently, the transmission risk for HAV, HEV and NoV GII via yellow and lesser mealworm and (tropical) house cricket to humans is low. However, other insect species and/or other foodborne viruses should still be investigated in future research, and it should involve industrial samples.

## Conclusions

In an extensive collection of industrially produced edible insects, this study uncovers a large contrast between the high risk for foodborne illnesses originating from bacterial spore formers and the low risk from human viruses in edible mealworms and crickets. For the spore-forming bacteria, the largest fraction of isolates was identified as members of the *Bacillus cereus* group, which contains foodborne and insect-borne pathogens. Sixty-five % of the *B. cereus* group isolates contained the cereulide plasmid that can allow the production of a heat-resistant toxin. The presence of the insect pathogens *Brevibacillus laterosporus* and *Lysinibacillus sp.*, on the other hand, may involve economic risks. Mitigation strategies include (i) the use of substrates that do not contain these species, which requires a thorough quality control, (ii) decontamination of the insects via the hurdle or combination strategy (e.g. combining heat with pressure, pulsed electric fields, irradiation, …) or (iii) prevent spores from germinating (e.g. by reducing the water activity, pH and/or temperature of the matrix, …). In literature, limited information is available, but insect producing and processing companies pay efforts in this field.

The presence of foodborne viruses in edible insects was investigated here for the first time. Using methods shown to be reliable, hepatitis A virus, hepatitis E virus and norovirus genogroup II were not detected (detection limit of 100 copies/µl extract). Hence, food safety risks related to these viruses can be stated to be very low. However, as industrial insect production methods are evolving due to automation and upscaling, virus detection is advised to be continued and also performed for other species than those considered in this study.

## Methods

### Sample collection and preparation

For experiment 1 (identification of bacterial endospores in edible insects), four samples of living yellow mealworms (*Tenebrio molitor*) and four samples of living house crickets (*Acheta domesticus*) were collected at the end of their rearing cycle. For each insect species, two different industrial rearing companies in Belgium (yellow mealworms) or Belgium and the Netherlands (house crickets) were sampled two times each (2 samples x 2 rearers x 2 sampling moments or batches). For experiment 2 (detection of foodborne viruses in edible insects), the same 17 mealworm and cricket samples that were investigated previously^7^ for their bacterial composition (yellow mealworm, house cricket and tropical house cricket (*Gryllodes sigillatus)* samples from different rearing companies and batches) were employed, as well as an extra sample of lesser mealworms (*Alphitobius diaperinus*) obtained in another study^4^ where it was used to study microbial dynamics during rearing (larvae day 35, post-harvest). After removal of dead specimens, insects were sedated by cooling them for approximately 1 h at 4 °C and subsequently, the samples were homogenised by pulverisation with a sterilised hand-held mixer (Bosch CNHR 25, Belgium) as described earlier^9^.

### Bacterial endospore counts and isolation

All samples employed in experiment 1 were subjected to aerobic and anaerobic bacterial endospore counts in threefold. Following the procedure described earlier^15^, for each analysis, 5 g of pulverised insects were diluted in 45 g peptone physiological salt solution (0.85% NaCl, 0.1% peptone, Biokar Diagnostics, Beauvais, France) and homogenised in a Bagmixer® (Interscience, Saint Nom, France) for 1 min. This primary dilution was given a pasteurisation treatment at 80 °C for 10 min and was then further diluted and plated on Plate Count Agar (PCA, Biokar diagnostics) using the spread plate technique to easily pick colonies later. Both aerobic and anaerobic endospore counts were determined after incubation for 48 h at 37 °C^48^. As a comparison, a similar but unpasteurised dilution series was used to determine the total viable aerobic and anaerobic counts of the samples (pour plate technique, PCA, 72 h at 30 °C). Anaerobic conditions were generated in Anaerocult containers (2.5 L, VWR International, Leuven, Belgium) using ‘AnaeroGen 2.5 L atmosphere generation systems’ (Thermo Fisher Scientific, Asse, Belgium) and evaluated using resazurin indicators (BR0055N, Thermo Fisher Scientific).

For each sample, after incubation, several colonies with various morphologies were picked from the countable plates and streaked on nutrient agar (NA, Biokar Diagnostics, 24 h at 37 °C, (an)aerobic incubation depending on their origin) to form axenic cultures. An individual colony was subsequently incubated overnight in nutrient broth (NB, Biokar Diagnostics, 18 h at 37 °C) and stored at - 80 °C after addition of glycerol to a final concentration of 50 % (v/v). This way, in total 67 and 95 spore-forming isolates were collected from the yellow mealworm and house cricket samples, respectively.

### Identification of bacterial endospores

All 162 endospore isolates were, after recultivation in NB, subjected to phenol/chlorophorm genomic DNA extraction as described earlier^49^ and subsequently to PCR (T100™ Thermal Cycler, Bio-Rad, Temse, Belgium), amplifying the 16 S ribosomal RNA (rRNA) gene. The PCR reaction (20 µl) contained 1.25 U of TaKaTa ExTaq Polymerase and 1 x ExTaq Buffer (Clontech Laboratories, Palo Alto, CA, USA), 312.5 µM of each dNTP, 1.0 µM of each primer (27F and 1492R, Table 3), and 5 ng genomic DNA (measured by a Nanodrop spectrophotometer, Thermo Fisher Scientific). PCR conditions included initial denaturation for 2 min at 95 °C, followed by 34 cycles of 45 s denaturation at 95 °C, 45 s annealing at 59 °C and 45 s elongation at 72 °C, and final elongation for 10 min at 72 °C. Obtained amplicons were sequenced using the same reverse primer as used in the PCR (1492R, Table 3) by Macrogen Europe (Amsterdam, the Netherlands). Resulting sequences were aligned and trimmed to an average read length of 861 bp. Isolates were subsequently classified using the EzBioCloud 16S rRNA gene database^50^. This way, a total of 142 isolates (50 from yellow mealworms, 92 from house crickets) could be identified.

**Table 3.**
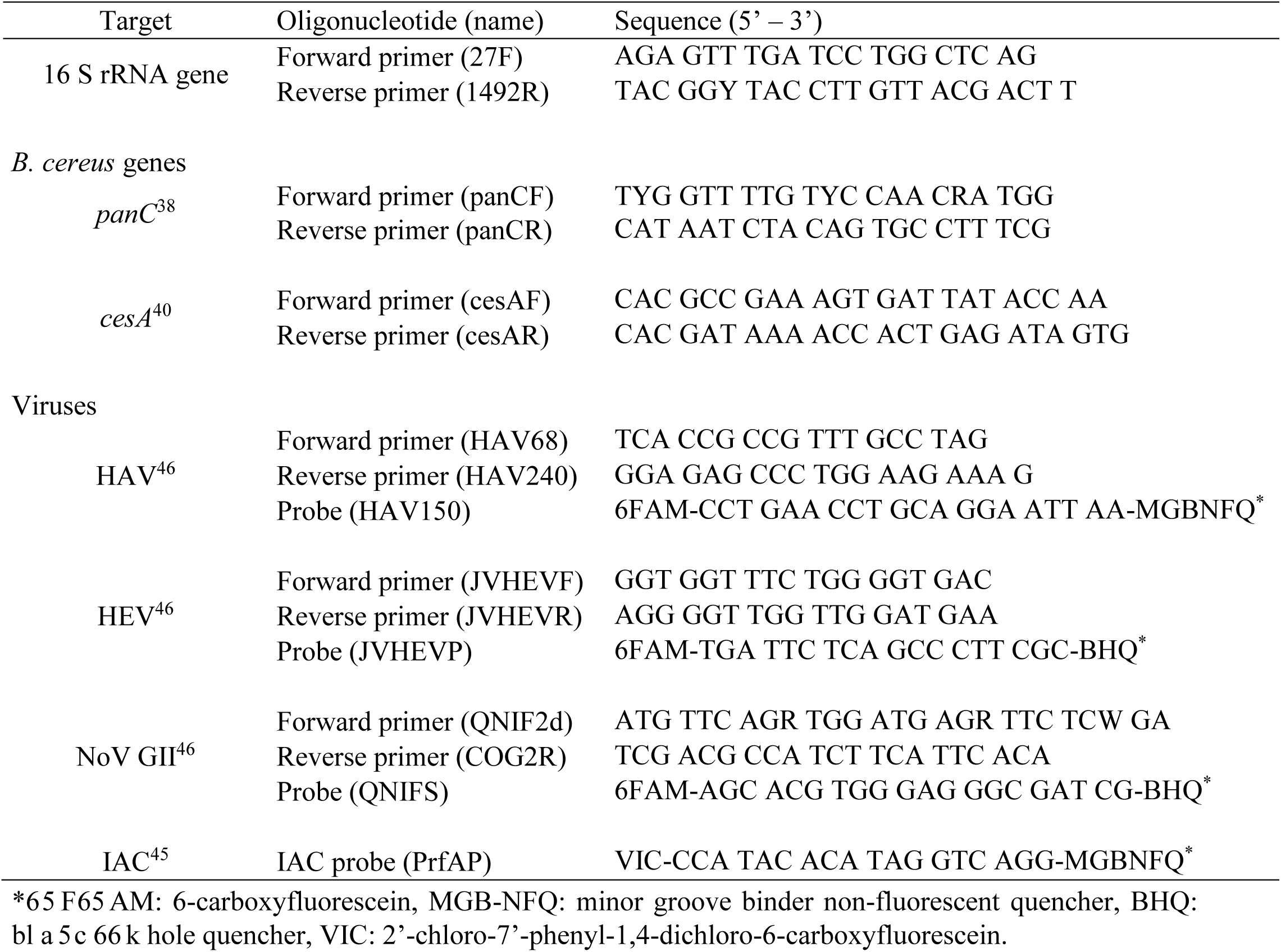
Primers and probes employed in this study.

### qPCR detection of B. cereus-related genes panC and cesA

For 99 isolates classified as *B. cereus* s.l., two additional genes were traced using qPCR. For each extract, firstly, a qPCR assay targeting the *panC* gene was applied in order to confirm the association with *B. cereus* s.l. Next, presence of the cereulide toxin pathway was investigated by assessing the *cesA* gene. All qPCR reactions (10 µl) were executed in a QuantStudio 3 qPCR system (Applied Biosystems, Thermo Fisher Scientific) and contained 5 µl PowerUp SYBR Green Master Mix with UNG (uracil-N glycosylase to prevent carryover contamination; Thermo Fisher Scientific), 1 µM (*panC*) or 0.4 µM (*cesA*) of each primer (Table 3) and 60 ng of DNA (measured by a MySpec spectrophotometer, VWR International). qPCR reaction conditions for the *panC* detection followed the master mix supplier’s protocol: 2 min at 50 °C (UNG activation) and 2 min of initial denaturation at 95 °C, followed by 40 cycles of 15 s denaturation at 95 °C and combined annealing and elongation for 1 min at 60 °C. For the *cesA* detection, the same conditions were applied, except for the initial denaturation step which was shortened to 20 seconds^51^. Each qPCR was followed by a melt curve analysis by raising the temperature gradually from 60 to 95 °C (0.1 °C/s) while continuously monitoring fluorescence. Positive (*B. cereus* emetic type, LMG 17603, Belgian Coordinated Collections of Microorganisms (BCCM)) and negative (no template, sterile ultra-pure water instead) controls were included. Data analysis was executed using the online application “Design and Analysis” in the qPCR DataConnect cloud software (Thermo Fisher Scientific). Successful detection was considered when the melt curve analysis revealed a single peak at the expected melting temperature, and when obtained Ct values did not exceed these of the negative control (33 for *cesA* and 35 for *panC*).

### RT-qPCR quantification of selected viruses

In order to detect and quantify HAV, HEV and NoV GII, previously described methods^46^ were optimised and used to screen the insect samples in experiment 2. Standard quantification curves and Internal Amplification Controls (IACs)^45^ were included to assess quantification and reliability. Corresponding virus reference cDNA was used as standard, either obtained from an RNA transcript (NoV GII after RT-PCR, see below) or plasmids containing virus genome fragments (HAV and HEV after PCR targeting the reference fragment). IAC RNAs were generated by transforming IAC-containing plasmids into chemically competent *Escherichia coli* cells (One Shot™ TOP10, Thermo Fisher Scientific) and transcription to RNA using the T7 High Yield Transcription kit (Thermo Fisher Scientific).

Samples were subjected in twofold to a total RNA extraction, using the Mo Bio RNA PowerSoil kit (manufacturer’s protocol, Carlsbad, CA, USA). Next, each RNA extract was subjected to a two-step reverse transcription real-time PCR (RT-qPCR) assay for each target virus separately. First, a tenfold RNA extract dilution was used to translate target virus RNA into cDNA (Table 3). In this step, the IAC corresponding to the target virus was added to exclude false negative results. Each reaction (20 µl) was performed using the High Capacity cDNA Reverse Transcription kit (Applied Biosystems, Thermo Fisher Scientific) and contained 2.0 µl RT Buffer, 1.0 µl MultiScribe™ Reverse Transcriptase, 2 µM reverse primer (Table 3), 0.8 µl dNTP Mix (100 mM), 0.5 µl RNase Inhibitor, 1000 IAC copies, and 40 ng RNA (as measured by MySpec spectrophotometer). Reaction conditions (iCycler Thermal Cycler, Bio-Rad) were as described by the supplier: 10 min at 25 °C, 2 h at 37 °C, and 5 min at 85 °C. Positive (10^8^ IAC copies) and negative (no template, sterile ultra-pure water instead) controls were included for all virus assays. Next, for each sample, the obtained cDNA was used as a template in the subsequent qPCR which aimed to detect and quantify HAV, HEV or NoV GII as well as to detect the corresponding IAC at the same time. The three virus assays were conducted based on protocols described earlier^46^. All analyses were performed in duplicate and each reaction (20 µl) contained 1x TaqMan Universal Master Mix with UNG (Applied Biosystems, Thermo Fisher Scientific), 0.25 µM of each primer (Table 3), 0.25 µM virus probe (FAM-labelled, Table 3) and 0.10 µM IAC probe (VIC-labelled, Table 3). All reactions were executed using a QuantStudio 3 qPCR system. qPCR conditions were as follows: 2 min at 50 °C, 10 min at 95 °C and 40 cycles of 15 s denaturation at 95 °C and 60 s combined annealing and elongation at 60 °C. Virus quantities and standard curve parameters were calculated in the qPCR DataConnect cloud software. Again, positive (1000 virus cDNA copies) and negative (no template, sterile ultra-pure water instead) controls were included. Only when the IAC was detected below the negative control Ct, the RT-qPCR assay was considered successful.

### Statistical analyses

To determine differences in microbial counts between samples in experiment 1, results were subjected to one-way ANOVA with Duncan post hoc test. Normality and homoscedasticity assumptions were tested for the analysed data sets using the Kolmogorov-Smirnov test and Levene’s test, respectively. SPSS Statistics 23 (IBM, New York, NY, USA) was used with significance level 0.05 for the statistical tests.

### Data availability

Sequencing data obtained in this study have been deposited in GenBank (National Center for Biotechnology Information, NCBI) with the accession codes MN508485 to MN508527 (Supplementary Table 1). All other relevant data are available from the corresponding author on request.

## Acknowledgements

Virus references and IACs were kindly provided by I. Di Bartolo from the Italian Istituto Superiore di Sanità (ISS, Rome, Italy) and N. Cook from Fera Science Ltd. (York, United Kingdom). Additionally, the authors would like to thank A. Paeleman (Scientia Terrae Research Institute, Sint-Katelijne-Waver, Belgium) for her expertise and assistance in designing and optimising the qPCR protocols and S. Crauwels (KU Leuven, Sint-Katelijne-Waver, Belgium) for processing the sequencing results. Also J. Franciotti, L. De Vrindt, N. Huybrechts, M. Gerits, S. Machtajiw, J. Plas and E. Van Vossole are acknowledged for their assistance in the lab. This research was financially supported by Flanders Innovation & Entrepreneurship (VLAIO) [Project 141129] as well as Internal Funds KU Leuven [PDM/18/159].

## Author Contributions

D.V. designed, prepared and executed all experiments, including sample collection and preparation, microbiological analyses, DNA and RNA extractions and PCR and RT-qPCR reactions. D.V. analysed the data, constructed tables and figures and wrote the main text. B.L. and L.V. supervised the study, provided additional insight in data analysis and revised the manuscript.

